# Association between chronic hepatitis C virus infection and myocardial infarction in people living with HIV in the United States

**DOI:** 10.1101/453860

**Authors:** Jessica Williams-Nguyen, Stephen E Hawes, Robin M Nance, Sara Lindström, Susan R Heckbert, H Nina Kim, W Chris Mathews, Edward R Cachay, Matt Budoff, Christopher B Hurt, Peter W Hunt, Elvin Geng, Richard D Moore, Michael J Mugavero, Inga Peter, Mari M Kitahata, Michael S Saag, Heidi M Crane, Joseph A Delaney

## Abstract

Hepatitis C virus (HCV) is common among people living with HIV (PLWH). The potential for extrahepatic manifestations of HCV, including myocardial infarction (MI), is a topic of active research. MI is classified into types, predominantly atheroembolic Type 1 MI (T1MI) and supply-demand mismatch Type 2 MI (T2MI). We examined the association between HCV and MI in the CFAR Network of Integrated Clinical Systems (CNICS), a multi-center clinical cohort of PLWH. MIs were centrally adjudicated and categorized by type using the Universal MI definition. We estimated the association between chronic HCV (RNA+) and time to MI adjusting for demographic characteristics, cardiovascular risk factors, clinical characteristics and substance use. Among 24,755 PLWH aged ≥18, there were 336 T1MI and 330 T2MI during a median of 4.2 years of follow-up. HCV was associated with a 68% greater risk of T2MI (adjusted hazard ratio (aHR) 1.68, 95% CI: 1.22, 2.30) but not T1MI (aHR 0.96, 95% CI: 0.63, 1.45). In a cause-specific analysis of T2MI, HCV was associated with a 2-fold greater risk of T2MI attributed to sepsis (aHR 2.26, 95% CI: 1.34, 3.81). Extrahepatic manifestations of HCV in this high-risk population are an important area for continued research.

---

Survival of people living with human immunodeficiency virus (HIV) (PLWH) has improved dramatically in the last 20 years, owing to a decline in mortality associated with antiretroviral therapy (1-5). During the same period, morbidity and mortality from non-AIDS-defining illnesses, including cardiovascular disease (CVD), have increased in this group (6-9). PLWH are at higher risk of myocardial infarction (MI) compared to the general population (10). Recent work has shown that among PLWH approximately half of incident MIs are Type 2 MI (T2MI) (11), whereas in the general population T2MI accounts for a much smaller proportion of MIs (12-20). In contrast to classical atheroembolic Type 1 MI (T1MI), T2MI results from myocardial oxygen demand-supply mismatch, a scenario that may result from a variety of conditions, including sepsis, illegal stimulant or drug-induced vasospasm, decompensated heart failure, and hypotension (21).

A large body of literature has linked CVD to chronic infection with hepatitis C virus (HCV), a common viral infection impacting an estimated 10 to 30% (22-26) of PLWH in the United States (US). Observational evidence supports an association between HCV and clinically evident CVD, in both the general population (27, 28) and in PLWH (29). Yet studies in the general population focusing on the role of HCV in coronary artery disease risk are inconclusive (27) and, when restricting to MI only, have generally not supported an association (30-33). One recent study found a higher risk of MI in those with HCV but only among the subgroup with high levels of total and low density lipoprotein (LDL) cholesterol (34). Among PLWH, very little research has assessed the association of HCV-coinfection with MI and, to our knowledge, none has examined whether such an association may differ by MI type.

Due to the higher prevalence of MI and HCV infection among PLWH, unclear associations between HCV and MI in the general population, different patterns of MI types among PLWH, and an increasing availability of highly effective treatments for HCV, it is important to establish whether HCV increases the risk of MI for PLWH. In this study, we sought to estimate the relative risk of incident T1MI and T2MI among PLWH who are coinfected with HCV compared to those who are not.

## Methods

### Study population

The Centers for AIDS Research (CFAR) Network of Integrated Clinical Systems (CNICS) cohort is a population of >32,000 PLWH who have received HIV care at one of 8 sites in the US from 1995 to the present and have consented to participation in CNICS research activities (35). Included in this study were all participants ≥18 years of age from six CNICS sites with comprehensive access to inpatient and outpatient electronic medical records (Johns Hopkins University; University of Alabama at Birmingham; University of California, San Diego; University of California, San Francisco; University of North Carolina at Chapel Hill; and University of Washington, Seattle). This analysis includes participants who contributed person-time during the years 1998 through 2016; the majority of follow-up occurred from 2010 to 2016. Entry into the study cohort was defined as six months after CNICS cohort entry or the first date of surveillance for MI events at the study site – whichever was later. Participants were followed until nine months after the date of last clinic visit or laboratory result (36), death, or the administrative censoring date for the site. Institutional review boards at each institution have approved CNICS research activities.

### CNICS data repository

Each CNICS site captures demographic, clinical, medication and laboratory data from all outpatient and inpatient encounters, including cardiac biomarkers, medications, diagnoses, and historical clinical information. In addition, participants were invited to complete an assessment of patient reported outcomes (PRO) and measures at 4-6 month intervals, capturing use of tobacco, alcohol and illicit substances as well as other health behaviors (37).

### Variables of interest

Ascertainment and adjudication of MI events in this cohort has been described in detail elsewhere (38). Briefly, identification of potential events was done centrally using multiple criteria, including MI-associated diagnostic codes (*International Classification of Diseases, Ninth Revision*, 410.00, 410.01, and 410.10), having an invasive cardiac procedure (percutaneous coronary intervention or bypass grafting), and elevated cardiac biomarkers. Sites then generated clinical data packets including primary data for all potential MI events. Packets contained chart notes, electrocardiograms, imaging and procedure reports, and laboratory values, but antiretroviral medication data were redacted. Packets were reviewed centrally by two physician experts to adjudicate MIs as definite or probable and to classify them as T1MI or T2MI, based on the Universal MI definition (11, 21, 38). While the Universal MI definition includes other types (e.g., cardiac procedure-related type 4 MI), these were uncommon (<10 cases) and so are not further discussed. In cases of discrepant findings, the assessment of a third reviewer was used to break ties (38).

Laboratory test results obtained in the course of routine care were used to determine HCV status. Chronic HCV infection was defined as a detectable result (i.e., above the lower limit of detection) for the most recent HCV ribonucleic acid (RNA) test before entry into the study cohort, and all remaining participants in the study cohort were classified as not chronically infected with HCV.

### Covariates

Covariates were taken from measurements available at study entry, except for height used to calculate body mass index which could be measured at any time. Diabetes was defined as any one of the following: hemoglobin A1c ≥ 6.5%, a clinical diagnosis of diabetes mellitus type 1 (T1DM) or type 2 (T2DM) and prescription of diabetes-related medication, or prescriptions of a diabetes-specific medication. Treated hypertension was defined as a clinical diagnosis of hypertension and documentation of an antihypertensive medication. Hepatitis B virus infection was defined on the basis of a positive result for hepatitis B e-antigen or surface antigen or deoxyribonucleic acid (DNA). Fibrosis-4 (FIB-4) index, a noninvasive measure of predicted liver fibrosis, was calculated as: [age (years) × aspartate aminotransferase (AST)]/[platelet count × alanine aminotransferase (ALT)^1/2^] (39) and categorized as >3.25 (severe fibrosis), 1.45-3.25 (indeterminate), <1.45 (no severe fibrosis).

History of injecting drug use was derived from HIV transmission risk factor reported at CNICS cohort entry and was coded as 1 if the participant reported membership in a transmission risk group that included injecting drug use. Ever-smoker was the presence of a tobacco-use diagnosis or self-report of tobacco use. Use of methamphetamine, cocaine/crack, marijuana, opiates (illicit use), cigarettes and alcohol at study cohort entry were available for a subset of participants (approximately 13%) as assessed by the ASSIST questionnaire (40). Drug and cigarette use categories were current, former or never use. Alcohol use was the AUDIT-C score derived from a short screening questionnaire about the frequency and quantity of alcohol consumed (41). Values range from 0 (least alcohol use) to 12 (most alcohol use).

### Statistical analysis

Approximately 87% of participants were missing information about methamphetamine, cocaine, opiate, marijuana, cigarette and alcohol use (from PROs) at the start follow-up. Missing values in all analytic variables were multiply imputed using chained equations with fully conditional specification in the R package mice (42) based on all other covariates yielding 100 complete datasets. For each complete dataset, the association between chronic HCV infection and time to the patient’s first MI event for each complete dataset was estimated using Cox proportional hazard regression models. Resulting inferences were pooled using Rubin’s Rules (43). Separate models examined T1MI, T2MI, cause-specific T2MI outcomes, and a composite outcome of all MIs. Models were adjusted for potential confounding factors. Minimally adjusted models considered only age and sex, while fully adjusted models included study site, demographic characteristics (age, sex, race/ethnicity, men who have sex with men), clinical characteristics (diabetes, treated hypertension, statin use, body mass index, lipid profile, lowest CD4+ cell count, hepatitis B virus infection, HIV viral load, antiretroviral therapy), history of injecting drug use, ever smoker and self-reported substance use (smoking, alcohol use, illicit substance use). Proportionality of hazards was assessed using Schoenfeld residuals for a random subset of imputations, and no consistent deviations were discovered. Statistical analyses were conducted in R version 3.4.3 (2017-11-30).

### Sensitivity analysis

Analyses were repeated to estimate the risk of MI outcomes associated with prior HCV infection in the absence of chronic infection at the time follow-up began. The exposed group for that analysis was those with a positive finding on the most recent HCV antibody test coupled with a negative finding for the most recent RNA test prior to baseline.

Because liver fibrosis was considered to be a potential mediator of the relationship between HCV and MI, it was not included in the regression models for the main analyses. However, in an effort to informally explore the extent to which liver fibrosis may serve as a mechanistic link between HCV and MI, the main analyses for T1MI, T2MI and all MIs were repeated with the stratum predicted to have severe fibrosis (FIB-4 > 3.25) and the stratum predicted to have no severe fibrosis (FIB-4 <1.45).

## Results

Among 23,407 PLWH, we observed 336 T1MI and 330 T2MI (666 MI total) during a median of 4.7 years of follow-up. In all, 2,280 (9.2%) had evidence of chronic HCV infection at the beginning of MI follow-up. The median age of participants was 40, 81% were male, and approximately half were non-white (Table 1). Compared with participants without HCV infection, those with HCV tended to be older and were less likely to report being men who have sex with men. Unsurprisingly, a much larger proportion (61.5% vs. 15.0%) had a history of injecting drug use (Table 1). The crude incidence rate for all MI events was 5.0 (95% CI: 4.7, 5.2) per 1000 person-years, and the rate was 4.6 (95% CI: 4.4, 4.9) among participants without chronic HCV and 8.2 (95% CI: 7.1, 9.4) among those with chronic HCV per 1000 person-years. T2MI events were attributed to a variety of causes (Table 2), with sepsis/bacteremia and stimulant-induced vasospasm being the most common.

**Table 1.**
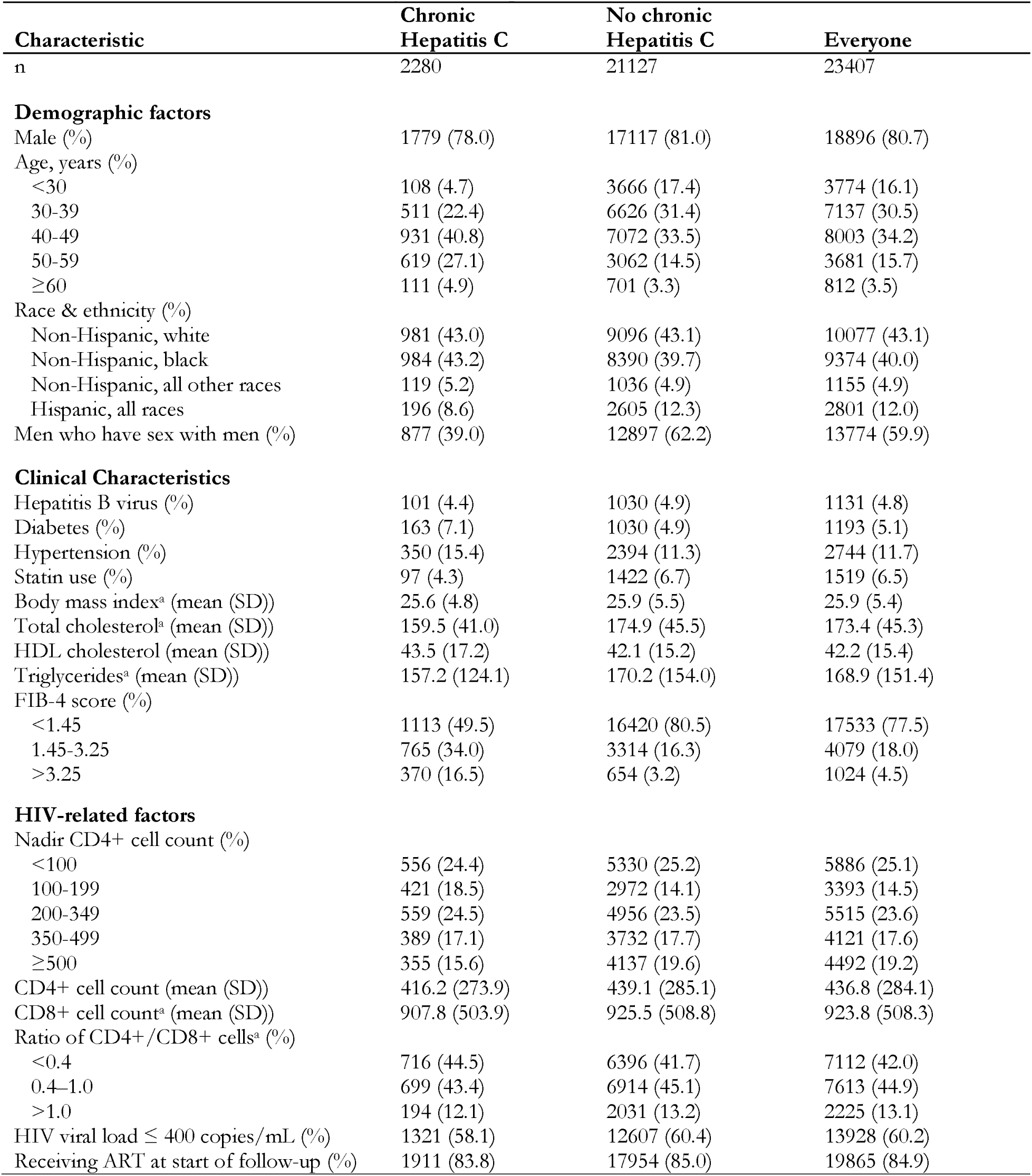
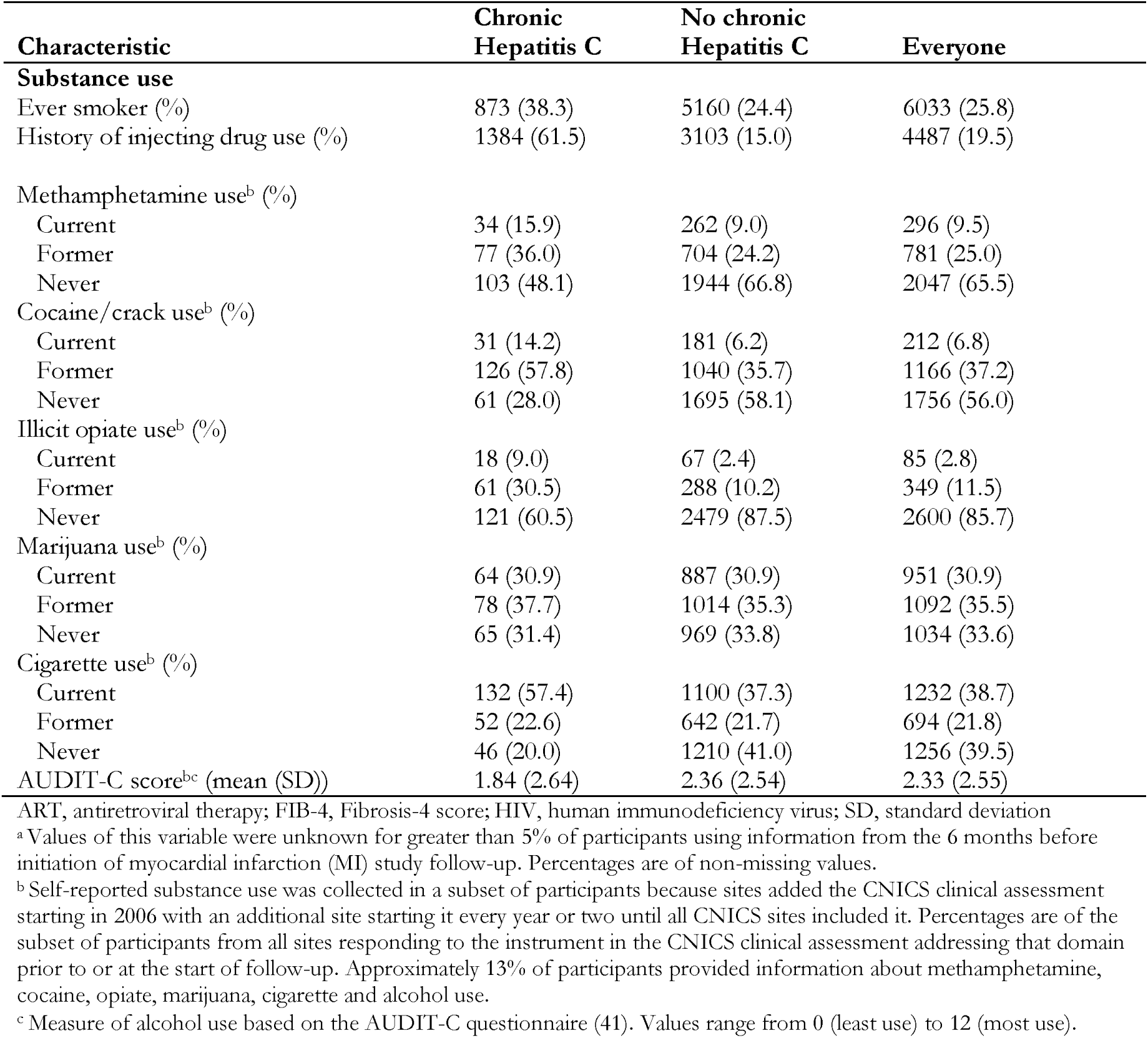
Baseline Clinical and Demographic Characteristics of Adults Living With HIV in Clinical Care at 6 CNICS Sites Across the U.S. in 1998-2016 by Chronic Hepatitis C Virus Infection Status

**Table 2.** Number of Type 2 Myocardial Infarctions (T2MI) Attributed to Causes Among Adults Living With HIV in Clinical Care at 6 CNICS Sites, 1998-2016

| **Attributed cause** | **Number of events (%)** |
| --- | --- |
| Sepsis/bacteremia | 118 (35.8) |
| Stimulant-induced vasospasm | 37 (11.2) |
| Hypertensive urgency/emergency | 35 (10.6) |
| Hypoxia | 24 (7.3) |
| Hypotension | 20 (6.1) |
| Arrhythmia | 17 (5.2) |
| Procedure related | 14 (4.2) |
| Gastrointestinal bleed | 11 (3.3) |
| Other rare causes | 54 (16.4) |
| **Total** | **330 (100.0)** |

Using a composite MI outcome, chronic HCV infection was associated with a 36% greater risk of MI after adjusting for demographic characteristics, traditional cardiovascular disease risk factors, clinical characteristics and substance use (adjusted hazard ratio (aHR) 1.36, 95% confidence interval (CI):1.06, 1.73) (Figure 1). Considering each MI type, chronic HCV infection was associated with a 68% higher risk of T2MI (aHR 1.68, 95% CI: 1.22, 2.30) after adjusting for potential confounders, but HCV was not associated with T1MI (aHR 0.96, 95% CI: 0.63, 1.45). In further analyses examining adjudicated causes of T2MI, HCV was found to be associated with an approximately 2-fold greater risk of T2MI attributed to sepsis (aHR 2.26, 95% CI: 1.34, 3.81) but was not associated with T2MI attributed to use of a stimulant (aHR 0.96, 95% CI: 0.15, 6.14) or other causes (aHR 1.47, 95% CI: 0.92, 2.34) after covariate adjustment (Figure 1). Because there were a small number of T2MI events attributed to use of a stimulant, we assessed the stability of the model for this outcome by estimating a model adjusted for age, sex, and history of injecting drug use – which gave relatively consistent results (Supplementary Table 1).

**Figure 1.**
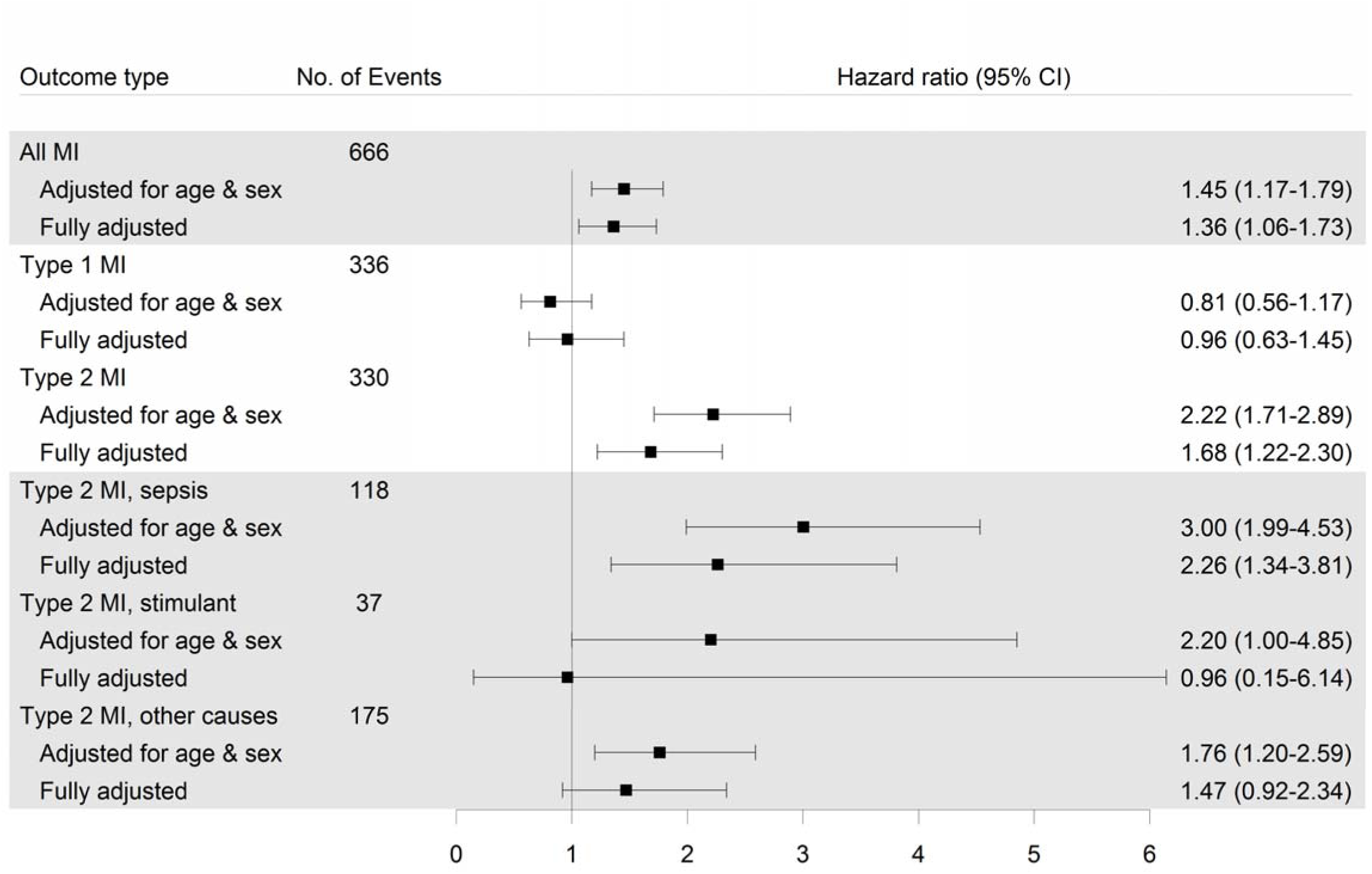
Forest plot of hazard ratios estimating the association of chronic hepatitis C virus infection with myocardial infarction outcomes among people living with HIV at 6 CNICS sites across the US. Fully adjusted models took into account age, sex at birth, race, ethnicity, site, men who have sex with men, ever smoker, history of injecting drug use, diabetes, statin use, hypertension, Hepatitis B virus, ART use, nadir CD4+ cell count, HIV viral load, body mass index, total cholesterol, HDL cholesterol, triglycerides, alcohol use score, and self-reported use of amphetamines, cocaine, opiates, marijuana, and cigarettes.

In sensitivity analysis, we investigated whether the risk of MI outcomes associated with chronic HCV appears to differ based on extent of liver fibrosis. In the stratum without predicted fibrosis (FIB-4 < 1.45), although HCV was associated with a 58% higher risk of T2MI after accounting for potential confounders (aHR 1.58, 95% CI: 0.95, 2.64), this relation was not statistically significant (Figure 2). In a separate analysis, we found that there was no association between prior HCV infection without chronic HCV infection (antibody-positive, RNA-negative) and any MI outcome (Supplementary Figure 1).

**Figure 2.**
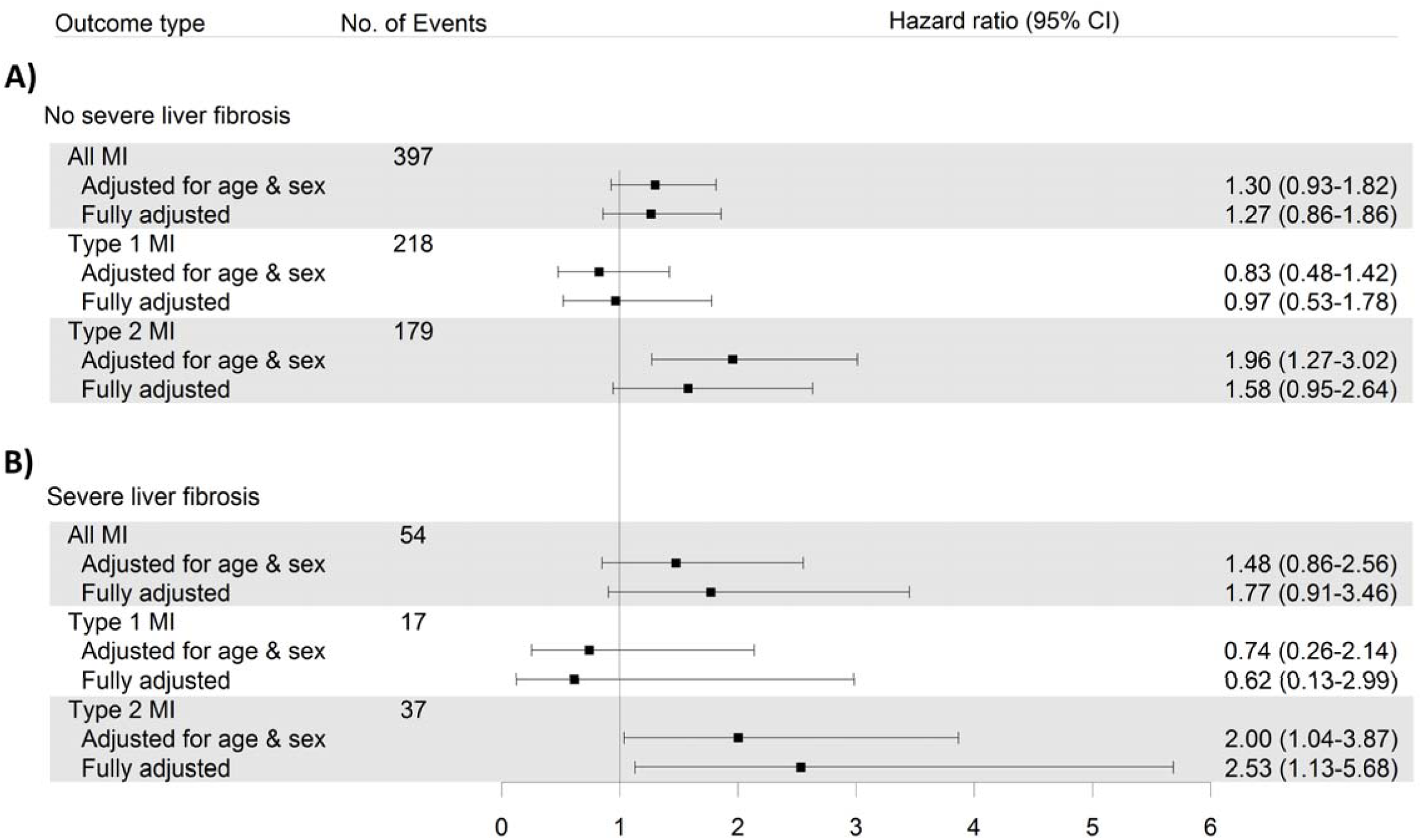
Forest plot of hazard ratios estimating the association of chronic hepatitis C virus infection with myocardial infarction outcomes among people living with HIV at 6 CNICS sites across the US in (**A**) the subset of participants with no predicted severe liver fibrosis (FIB-4 score < 1.45) and (**B**) the subset with predicted severe liver fibrosis (FIB-4 score > 3.25). Fully adjusted models took into account age, sex at birth, race, ethnicity, site, men who have sex with men, ever smoker, history of injecting drug use, diabetes, statin use, hypertension, Hepatitis B virus, ART use, nadir CD4+ cell count, HIV viral load, body mass index, total cholesterol, HDL cholesterol, triglycerides, alcohol use score, and self-reported use of amphetamines, cocaine, opiates, marijuana, and cigarettes.

## Discussion

In this cohort of PLWH across the US, we found that HCV/HIV-coinfected persons experienced a greater risk of T2MI and this was particularly driven by the relation between chronic HCV infection and T2MI attributed to sepsis. In contrast, there was no significant increase in the risk of having T1MI among coinfected persons. To our knowledge, this is the first study to assess the association between HCV infection and MI types, particularly T2MI, among PLWH.

In general, it is our findings with respect to T1MI that are likely to be most comparable to prior studies of this question, which in the general population have not found an association between chronic HCV infection and MI (30-33). Two prior studies in PLWH were inconclusive. Bedimo et al. (44) reported a 25% higher risk of acute MI in U.S. veterans infected with HCV and HIV compared to those infected with HIV alone, however this association did not reach the level of statistical significance, while the D:A:D study group found no association between HCV seropositivity and MI (45). Although both of these studies ascertained composite MIs without reference to type, both defined MIs using administrative diagnosis codes. The proportion of MIs detected in prior studies that were T1MI versus T2MI is not known, however the ability of administrative codes to detect T2MI seems generally poor (46). Therefore, prior evidence appears to concur with our finding that chronic HCV is not associated with classical, atheroembolic T1MI.

We observed a greater risk of T2MI associated with HCV and, more specifically, an approximately 2-fold greater risk of T2MI due to sepsis in those coinfected with HCV and HIV compared to those infected with HIV alone. This may be related to consequences of HCV infection, such as liver dysfunction and/or chronic inflammation, or differences in sepsis risk factors between HCV-infected and uninfected participants. While our results suggest a higher risk of T2MI attributed to sepsis in those with evidence of chronic HCV infection, we did not detect a similar relation when considering prior HCV infection without evidence of viremia. This lends credence to the hypothesis that ongoing chronic infection or associated biological factors (in addition to risk behaviors associated with HCV exposure) may play a role in the risk of T2MI resulting from sepsis or of sepsis, itself. A large study of hemodialysis patients in the US, which found that the risk of bacteremia was greater in those with HCV infection, provides support for HCV as a risk factor for sepsis (47), and the landmark REVEAL study found that HCV seropositivity is associated with an 50% higher risk of mortality due to circulatory diseases (48).

In sensitivity analysis, stratification by predicted liver fibrosis score (FIB-4) suggested potential heterogeneity in the association of HCV with T2MI by extent of liver fibrosis. In those without predicted fibrosis, chronic HCV was not significantly associated with T2MI after accounting for potential confounders. Those with predicted severe fibrosis were uncommon in this cohort, and there were few MI events in that subgroup. Although we have presented the estimated relative hazard of MI outcomes among those with severe fibrosis, we are limited in our ability to discern whether risk of T2MI associated with HCV is greater in this subgroup as small sample size may have introduced instability into the relative risk estimates. In addition to overall replication of our findings, an ideal future study in this area would more comprehensively investigate the role of liver fibrosis in the relation of HCV with cardiovascular disease outcomes, particularly in those with advanced fibrosis. Bacterial infections and sepsis are known complications and major sources of morbidity for patients with cirrhosis (49).

Strengths of this study include adjudicated ascertainment of MI events including typing, extensive follow-up of participants providing a wide range of information about potential confounders, and the CNICS PRO assessment with the use of multiple imputation methodology that permitted inclusion of self-reported substance use in our analytic models. HCV infection status in most studies of HCV and MI was defined on the basis of either seropositivity for anti-HCV antibodies or administrative diagnosis codes for HCV (30-33, 44, 45). These measures are likely to be less specific for chronic HCV infection compared to ascertainment on the basis of an RNA test as we did here. One study of the general population in an Arkansas medical system found that participants with detectable HCV RNA had a greater incidence of coronary heart disease than those who tested positive for antibodies against HCV but without detectable HCV RNA. In combination with the findings presented here, this supports the use of HCV RNA testing to identify chronic HCV, something that should be highly feasible with the increasing uptake of nucleic acid based tests for HCV in clinical practice (50).

We were unable to differentiate whether HCV is associated with the incidence of sepsis itself, or T2MI as a complication of sepsis. This is an important area for future work. Because adjudication of causes of T2MI events must necessarily rely on expert judgement, some measurement error may have occurred when categorizing these complex medical events. Our study was also not sufficiently powered to assess causes of T2MI other than sepsis and stimulant-induced vasospasm. We also used HCV RNA testing performed in the course of routine clinical care, and therefore potentially missed some cases of chronic HCV in those who did not undergo RNA testing in that setting. Finally, because any study seeking to characterize the effects of HCV infection must be observational, the possibility of unmeasured and residual confounding cannot be completely excluded. In particular, we note that 87% of participants were lacking baseline information for self-reported substance use and time-varying substance use was not included in this analysis, leaving open the possibility of confounding by differences in substance use in those with and without chronic HCV. However, this rich database of clinical and patient-reported information, combined with analytic techniques accommodating missingness in covariates, enabled us to adjust for many important potential confounders.

In this large and diverse multicenter cohort of PLWH in the US, we found that chronic HCV infection at the beginning of follow-up was not associated with incident classical atheroembolic T1MI but was associated with T2MI attributed to sepsis. These findings demonstrate the importance of examining MIs by type among PLWH and support increasing calls for broad-scale efforts to better understand the biological underpinnings of T2MI, and ultimately identify effective management approaches (51). Further research is needed to replicate these findings, elucidate biological mechanisms and determine whether risk of sepsis and/or T2MI might be an important consideration for HCV treatment decisions in this high-burden population.

## Supplementary material – Williams-Nguyen, et al.

**Supplementary Figure 1.**
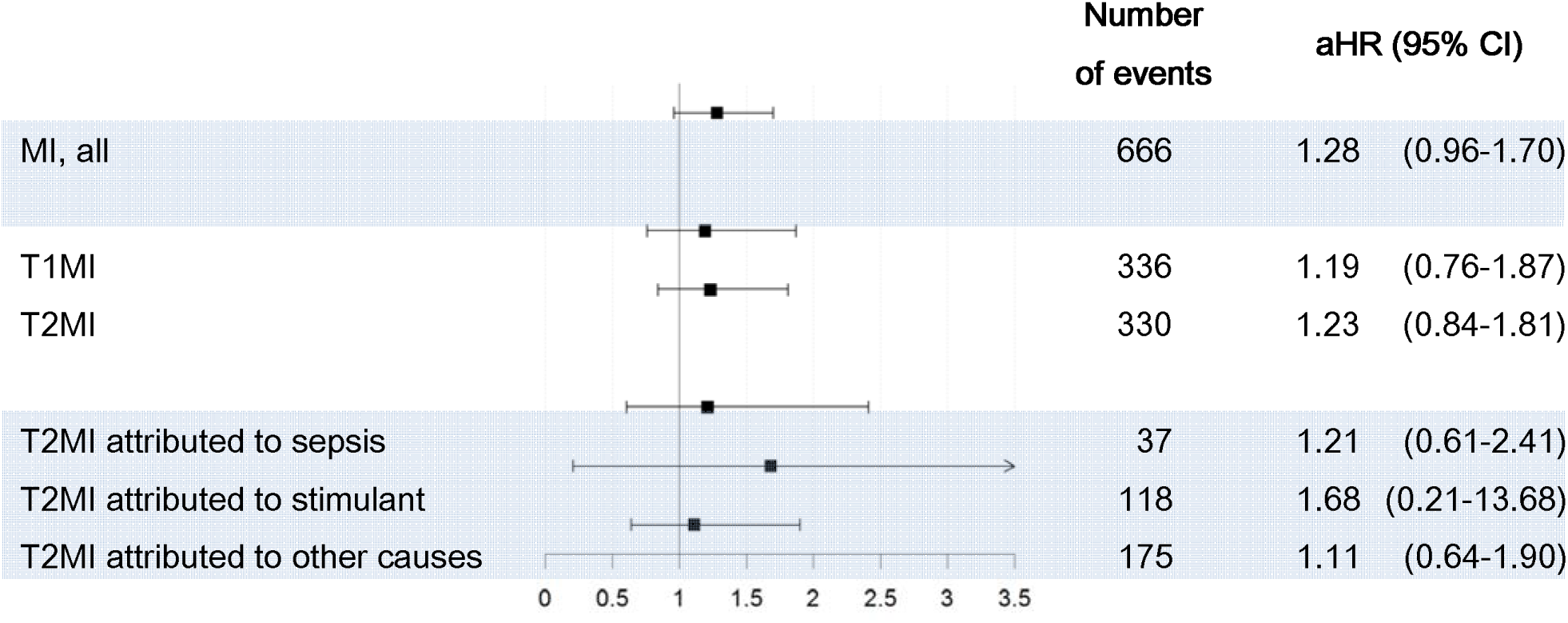
Forest plot of adjusted hazard ratios for incident MI outcomes in those with prior HCV infection without chronic infection compared to those without evidence of prior HCV infection (antibody negative). Prior HCV infection without active infection at the beginning of MI follow-up was defined as a positive result on the most recent HCV antibody test combined with a negative result on the most recent HCV RNA test. Hazard ratios are from Cox proportional hazard regression models adjusted for age, sex at birth, race, ethnicity, site, men who have sex with men, ever smoker, history of injecting drug use, diabetes, statin use, hypertension, Hepatitis B virus, ART use, lowest CD4+ cell count, HIV viral load, body mass index, total cholesterol, HDL cholesterol, triglycerides, alcohol use score and self-reported use of amphetamines, cocaine, opiates, marijuana, and cigarettes. Multiple imputation was used and parameters were pooled using Rubin’s rules. Abbreviations: aHR, adjusted hazard ratio; CI, confidence interval; MI, myocardial infarction; T1MI, Type 1 MI; T2MI, Type 2 MI

**Supplementary Table 1.** Hazard ratios describing the association of chronic HCV with T2MI attributed to stimulant use

| Adjustment variables | Hazard ratio | 95% Confidence interval |
| --- | --- | --- |
| None (crude estimate) | **2.60** | **1.19, 5.70** |
| Age, sex | 2.20 | 1.00, 4.85 |
| Age, sex, history of injecting drug use | 1.18 | 0.51, 2.75 |
| All covariates without substance use PROs | 1.07 | 0.45, 2.55 |
| All covariates with substance use PROs | 0.96 | 0.15, 6.14 |

Hazard ratios are from Cox proportional hazards regression models estimating incident myocardial infarction outcomes in those with chronic HVC infection compared to those without evidence of chronic HCV infection. Covariates were age, sex at birth, race, ethnicity, site, men who have sex with men, ever smoker, history of injecting drug use, diabetes, age, sex at birth, race, ethnicity, site, men who have sex with men, ever smoker, history of injecting drug use, diabetes, statin use, hypertension, Hepatitis B virus, ART use, lowest CD4+ cell count, HIV viral load, body mass index, total cholesterol, HDL cholesterol,
triglycerides, alcohol use score and self-reported use of amphetamines, cocaine, opiates, marijuana, and cigarettes. Multiple imputation was used and parameters were pooled using Rubin's rules. **Boldface** indicates significance at $\alpha=0.05$ .

## Abbreviations

aHR,: adjusted hazard ratio
AIDS,: acquired immunodeficiency syndrome
ART,: antiretroviral therapy
CFAR,: Centers for AIDS Research
CI,: confidence interval
CNICS,: CFAR Network of Integrated Clinical Systems
CVD,: cardiovascular disease
FIB-4,: Fibrosis-4 score
HCV,: hepatitis C virus
HIV,: human immunodeficiency virus
LDL,: high density lipoprotein
MI,: myocardial infarction
PLWH,: people living with HIV
RNA,: ribonucleic acid
T1MI,: Type 1 myocardial infarction
T2MI,: Type 2 myocardial infarction

